# nanoDoc: RNA modification detection using Nanopore raw reads with Deep One-Class Classification

**DOI:** 10.1101/2020.09.13.295089

**Authors:** Hiroki Ueda

## Abstract

Advances in Nanopore single-molecule direct RNA sequencing (DRS) have presented the possibility of detecting comprehensive post-transcriptional modifications (PTMs) as an alternative to experimental approaches combined with high-throughput sequencing. It has been shown that the DRS method can detect the change in the raw electric current signal of a PTM; however, the accuracy and reliability still require improvement. Here, I present a new software program, named as nanoDoc, for detecting PTMs from DRS data using a deep neural network. Current signal deviations caused by PTMs are analyzed via Deep One-Class Classification with a convolutional neural network. Using a ribosomal RNA dataset, the software archive displayed an area under the curve (AUC) accuracy of 0.96 for detecting 23 different types of modifications in *Escherichia coli* and *Saccharomyces cerevisiae*. Furthermore, I demonstrated a tentative classification of PTMs using unsupervised clustering. Finally, I applied this software to severe acute respiratory syndrome coronavirus 2 data and identified commonly modified sites among three groups. nanoDoc is an open source software (GPLv3) available at https://github.com/uedaLabR/nanoDoc

**Author Summary:** RNA post-transcriptional modifications (PTMs) is regulate multiple aspects of RNA function, including alternative splicing, export, stability, and translation, and the method to identify multiple types of PTMs is required for further advancement of this fields called ‘epitranscriptomics’. Nanopore singlemolecule direct RNA sequencing (DRS) can detect such PTMs, however the accuracy of the method needs to be improved. Detecting PTMs can be solved as a One-Class Classification problem, which is widely used in machine learning fields. Thus, a novel software named ‘nanoDoc’ for detecting PTMs was developed. The nanoDoc use convolutional neural network to extract the feature signal from nanopore sequencer and Deep One-Class Classification to detect PTMs as an anomaly. The software archive displayed an area under the curve (AUC) accuracy of 0.96 for detecting 23 different types of modifications in *Escherichia coli* and *Saccharomyces cerevisiae.* This software is applicable to different samples, and tested on severe acute respiratory syndrome coronavirus 2, and human transcript data as well.

## Introduction

To date, over 150 different types of post-transcriptional modifications (PTMs) have been identified. PTMs have long been known, in tRNA and rRNA, play important roles in maintaining structure and adjusting codon usage. More recently, an increasing number of PTMs have been identified in other types of RNA (e.g. mRNA and non-coding RNA), which regulate multiple aspects of RNA function, including alternative splicing, export, stability, and translation. PTMs play a role in the life cycle of various viruses and in the cellular response to viral infection [1]. One such PTM is N^6^-methyl adenine (m6A), introduced by the adenine methyltransferase family of enzymes (writers), which is specifically recognized by a set of RNA-binding proteins (readers). Additionally, studies have shown that fat mass and obesity-associated protein (FTO) and ALKBH5 exhibit m6A demethylase (eraser) activity [2][41]; hence, PTMs are dynamic. The concept of the activity balance between the writers, readers, and eraser in modulating cellular functions is the basis for the rapidly developing field of “epitranscriptomics”. Therefore, transcriptome-wide PTM identification is a key technology for epitranscriptomics.

For the first generation of epitranscriptomics tools, experimental methods, including detection of reverse transcription error patterns (RT-signature) with/without a specific chemical [3][4][5][6][7] and immunoprecipitation using antibodies specific to certain PTMs, [8][9], have been developed. Although these experimental methods have been adapted for the different types of PTMs, they also have limitations. For example, with respect to RT-signature detection, chemical conditions are available for only a few types of modifications, and it is difficult to detect all modification sites if they are located in close proximity. Likewise, immunoprecipitation tends to exhibit problems related to resolution and antibody specificity. Moreover, these experimental methods can only detect one type of modification; as different types of modifications are dynamically acting in cells, a comprehensive method for detecting multiple PTMs, with a simple setup, is necessary for further advancement of epitranscriptomics.

Nanopore single-molecule direct RNA sequencing (DRS) can detect PTMs in addition to performing nucleotide base calling [10][11]. DRS has the advantage of direct detection of RNA molecule, in contrast to short-read sequencing, in which information of RNA modification is lost via the process of reverse transcription and polymerase chain reaction. To detect DNA modifications, recurrent neural network (RNN) models, such as DeepSignal [12] or DeepMod [13], have been developed. However, there are numerous types of RNA modifications (e.g. at least 19 types in mRNA), making the application of RNN models difficult, in terms of deep neural network model building and preparing the training data set. One approach for acquiring the training set for RNA modification is via chemical synthesis of modified bases [14]. However, chemical synthesis is only available for selected PTMs, and it is difficult to generate all combinations of the modified and unmodified states of the nucleotide sequence. Therefore, most current PTM detection algorithms rely on base-calling errors induced by modifications and external information, such as modification motifs or previously verified modification sites [15] [14]. However, these approaches rely on external errors of unknown processing, and that any signal may be base-called as a same base to the reference. In addition, using a priori data, such as modification motifs, for filtering is problematic when the data source contains errors, and any filtering method may lead to bias that suppresses de novo discovery.

In another type of algorithm, the raw current signal is aligned against a theoretical signal by either the Viterbi algorithm with the hidden Markov model (HMM) [16] [17] or RNN [18], or adaptive banded dynamic time warping [19] and the difference in raw signal levels is compared. As the current signal is disturbed by the existence of PTMs, statistical testing such as the Kolmogorov–Smirnov (KS) test [19][20], is used to detect PTMs. Identifying modification sites with these methods is useful and may lead to further classification of different PTMs at detection sites; however, the accuracy of these methods must be improved.

Detecting PTMs from non-modified *in vitro* signaling can be solved by One-Class Classification, which is widely used in machine learning. For example, Perera et al. [21] developed a state-of-art One-Class Classification algorithm, Deep One-Class (DOC), with a convolutional neural network (CNN). Recently, CNN has been used to analyze of nanopore data; for example, DeepBinner [22] can detect barcode sequences more accurately than conventional base caller and mapping reads approaches. Thus, using the DOC and CNN algorithms, I present a novel software for detecting PTMs.

## Method

### nanoDoc Design overview

The program has two parts: the training part and inference (PTM detection) part. The training parts involve two processes, initial training and DOC training. In initial training, one-dimensional (1D)-CNN is pretrained to learn the general futures of the input signal for 1024 5-mers (Fig 1b). In the next DOC, the learned weight in the initial training is used for the lower layer of the neural network (transfer learning) and DOC learning to obtain a specific future to distinguish the target 5-mer to a PTM as an anormal signal (Fig 1c). In the inference (PTM detection) part, the learned neural network is applied to a raw current signal of wild-type RNA (Fig 1d and 1e).

**Fig 1.**
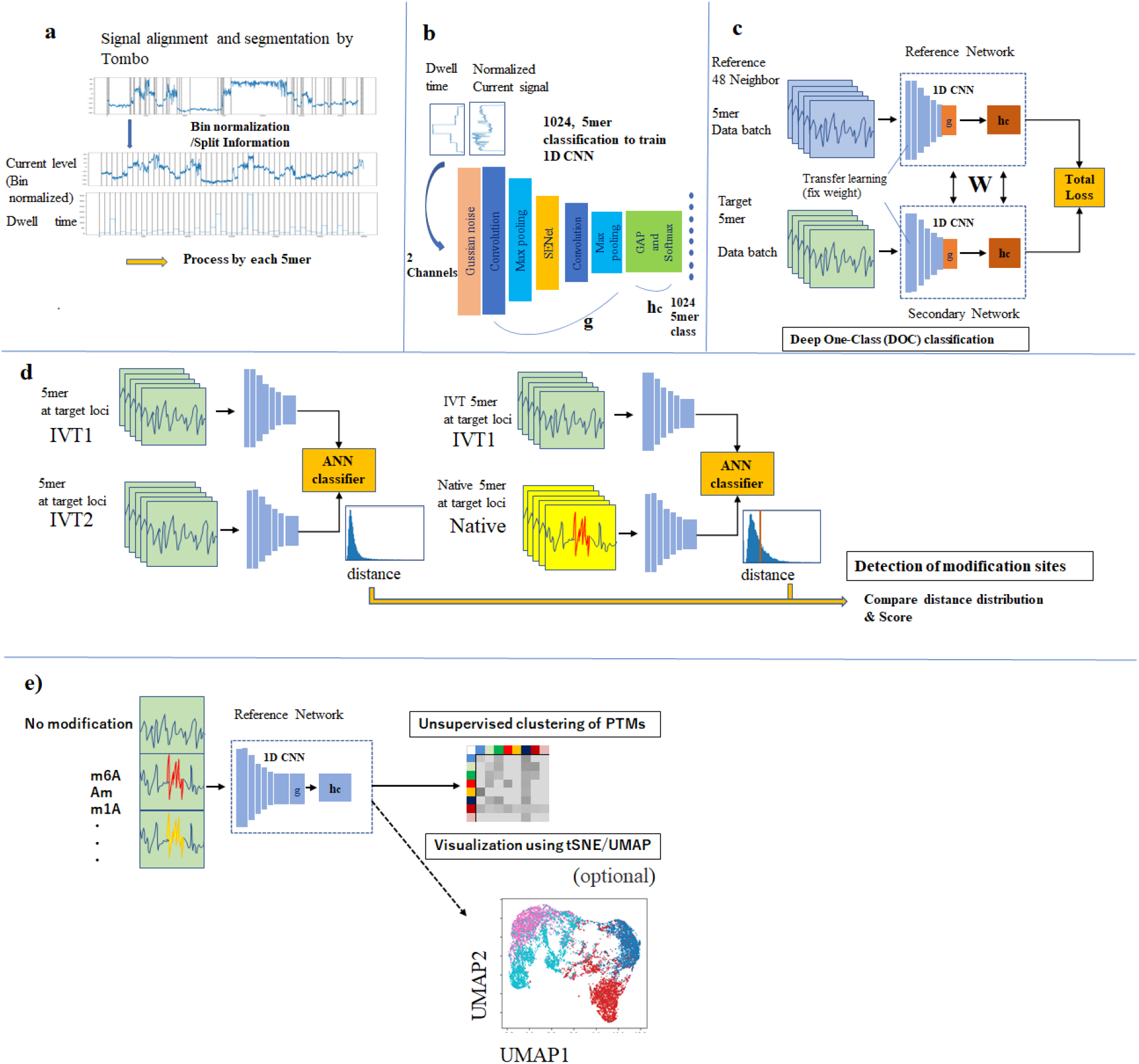
Modification detection algorithm by nanoDoc. The modification detection algorithm by nanoDoc a) Decomposition of resquiggled signal to uniform length of 5-mer. The resquiggled signal was decomposed into two channels: one for current level and the other for dwell time interval. The current was either up- or down-sampled to fit in the uniform length interval. b) ) 1D-CNN was trained with 1024 labeled 5-mer combinations. c) Deep One-Class (DOC) training for each 5-mer. (modified from *Learning Deep Features for One-Class Classification*[21]) Reference 5-mer and 5-mer with similar sequences were subjected to DOC classification. d) Scoring with approximate nearest neighbor Euclidian distance. Score is assigned based on Euclidian distance to the reference dataset. e) Unsupervised clustering of multiple PTMs. The feature 5mer data were subject to k-mean unsupervised clustering

### Iinitial training of 1D-CNN

From the *in vitro* (IVT) sequence of previous studies [27][14], the 5-mer signal was extracted for all 1024 possible 5-mer combinations for initial CNN training. The CNN was constructed using Keras [28] with Tensorflow [29] backend. A 1D-CNN was constructed to capture the specific features of the Nanopore signals. The CNN architecture consisted of 64 layers, including convolutional, batch normalization, and drop out layers reinforced by Squeezenet [30] architecture (S2 Fig). nanoDoc used two channels of data as inputs: the first channel was the normalized current signal of the 5-mer, and the second channel was dwell time information formatted to the same bin size. Class classification of the 1024 5-mer combinations was performed using 1,000 signals for training and 200 signals for validation (Fig. 1b). Four NVIDIA Tesla v100 graphics processing units were used, and training was conducted with up to 500 epochs for 454 min. The epoch of 340 yielded the best validation accuracy of 0.32 (S3a Fig) Several other models were tested when choosing this specific model (S13 Fig).

### Deep One-Class learning

DOC classification was performed using the trained CNN. First, the weight with best accuracy (the epoch of 340) from the previous procedure was loaded into the CNN model. The top three layers were removed, and the convolutional layers (g) were connected in a parallel manner with an additional sub classification network layer (hc), where the weight was shared between two CNNs (Fig. 1c). Each CNN was subject to transfer learning, where the weight was fixed, except for the top six layers (Fig. 1c).

Next, DOC classification was performed for each of the 1024 5-mer combinations. The 5-mer raw data were utilized as input for the secondary CNN. Reference data were selected from 48 other 5-mers with nucleotide sequences similar to the target (S4a Fig.); the loss function and how to choose neighbor sequences). The networks were trained for 20 epochs (S4b Fig); however, the final accuracy was decreased after epoch 2, possibly because of overfitting as indicated in the DOC paper [21]. Thus, the weight of epoch 2 was used for all 5-mers. For the training and test, 12,000 signals were extracted for each kmer sequence from the IVT unmodified sample, 10,000 signals were used for training, and 2000 signals were used for testing. The dataset used in training is summarized in S4 Table.

### Inference and distance search

At each genomic locus, IVT sequences, twice as many as the native sequences, were selected for detection, to create an equal number of sequences in three raw data batches: IVT1, IVT2, and native. A maximum of 20,000 signals was utilized for each data batch. For each 5-mer at the genomic locus, the DOC-trained weight was loaded to the CNN, along with three data batches, as the program progressed along the reference. In the output layer, the 320 × 2 input dimension was reduced to 16 dimensions. The Euclidian distances of the output vectors, from IVT1 to IVT2 and IVT1 to native, were compared using approximate nearest neighbors (ANN)[31] with Faiss [32]. For each entry vector, five of the nearest vectors in the target dataset were selected, and the mean distance was summarized as the distribution (Fig. 1d).

### Scoring

The Euclidian distance from the native signal to the IVT signal were compared and normalized for scoring. (S1 methods).

### Clustering and extracting the feature

The 16-dimensional feature output of the native sequence was subjected to k-mean clustering and dimensional reduction uniform manifold approximation and projection (UMAP) [33].

## Results

### Detection of known PTMs in rRNA and evaluation

The nanoDoc has been applied to the native rRNA sequencing data of *Escherichia coli* and *Saccharomyces cerevisiae* from a previous study[20]. The data was base-called and processed using Tombo, as described in the previous section, using an appropriate reference sequence (S2 Table). In addition, the list of known modification sites and KSStats values (generated by Tombo in comparative mode) were downloaded from the GitHub repository of nanoShape [20]. The score distributions across the genomic position of each rRNA are depicted in Fig 2a, where the red line represents the positions of known PTMs and blue line shows the scores. For accuracy evaluation, I conducted receiver operating characteristic (ROC) analysis with nanoDoc, KSStats, and mean current differences from Tombo. The results are shown in Fig 2b, with the ROC curves for nanoDoc (blue), KSStats (Stephenson et al; green), and Tombo mean current difference (orange). The area under the curve (AUC) values were 0.96, 0.89, and 0.79, respectively (S2 Method). We examined the nanoDoc score for each of the 23 modification types (Fig 2c). All scores at the 147 modification sites were higher than the 75^th^ percentile baseline, except for one m5U modification site. The baseline was drawn from the collection with modification sites. Note that the actual modification frequency at each known modification site was unknown, and a low score may represent the low modification frequency at a given position. Pseudouridine (Ψ) and Nm (2’-O-methylation) showed the highest scores (0.19±0.16 out of 53 sites and 0.26±0.19 out of 57, respectively) among the PTMs, which implies that some modified bases and modifications on ribose are prone to detection. Although the sample sizes for some other modification sites were not sufficient for general evaluation, the overall result suggests that majority of base modifications could be detected using this method.

**Fig 2.**
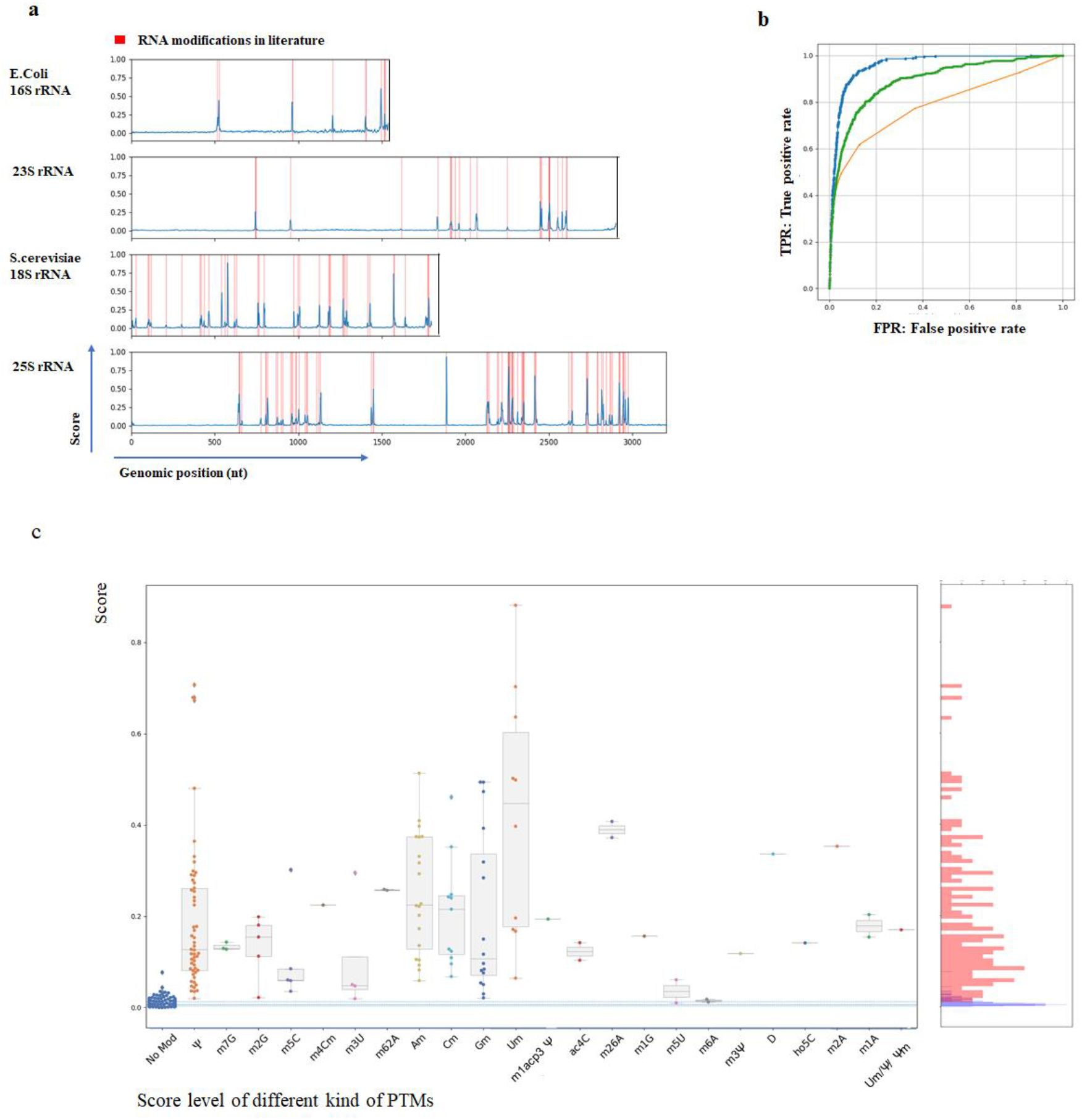
Evaluation of nanoDoc accuracy. a) Evaluation of nanoDoc using ribosomal RNA of *E.coli* and S. *cerevisiae*. The red line show the position of known modification sites in literature. The bule line show the score level of the nanoDOC at each position b) ROC curve nonoDoc (blue), KS statistics Stephenson et.al. (green) and Tombo mean current difference (orange) AUC values are 0.96,0.89,and 0.79 respectively c) Score levels are summarized by each modification type, and baseline shows data with no modification (left most). Percentile lines for base lines were drawn at 75%,50%, and 25% from the top. The right panel show the score distribution of modified (red) and unmodified (blue) sites.

For m6A modification, the score level was low (0.02 ± 0.004) in the rRNA dataset. To investigate m6A detection more precisely, I analyzed synthesized RNA data from a previous study [14]. I examined 81 types of 5-mers with m6A at the center, “BB (m6A) BB” (where B = G, C, or U), against the reference “BB (A) BB” 5-mer. The AUC value for detecting m6A modifications, in this dataset, was 0.78 (S6 Fig). For the DRACH motif, the AUC value was 0.88. (S6 Fig) in the modified dataset. Since all adenosines were chemically modified to m6A, which may have affected the score of the unmodified 5-mer neighboring 5-mer.

Although a small AUC value suggests that the method must be improved to accurately detect m6A modifications, the m6A modification score was mostly within the detection range in the known m6A motif (DRACH motif). According to 3-mer analysis, G(m6A)C, C(m6A)G or G(m6A)U, and U(m6A)G are prone to detection compared with other sequence combinations. (S6 Fig)

I also examined how the modification frequency correlates with the score, using the frequency data from a previous study [34] (Supplementary Fig. 7,12). The results showed no correlation (r^2^ = 0.15) between the score and the previously reported modification frequency. Additionally, the data of depth analysis indicated that the modification types were more closely related to the score value than the modification frequency. The read depth also affects the score, even after recalibration with the depth factor (S7 Fig). The relative score intensities showed similar trends among groups, suggesting that the modification type is the strongest factor affecting the final score.

The possible reason for this low correlation is that the magnitude of current disturbance depends on the background 5-mer sequence composition, making it difficult to estimate the modification frequency from the nanoDoc score alone. Thus, further classification and data accumulation are necessary to predict the modification frequency from the score.

### Clustering analysis of 5-mers with multiple modifications

Within the test data set, seven types of 5-mers have multiple PTMs at different positions (S1 Table). For those seven types of 5-mers, raw signals were applied to nanoDoc CNN and the future value was extracted (Fig 1e) into a 16-dimensional vector. The output vector was subjected to k-means clustering (Fig 3). The figure in the top row depicts the UMAP representation of the extracted vector (S3 Method), where data from different positions is colored differently. The second row is the result of k-means clustering, with data in the same cluster given the same color. Note that clustering was performed in a 16-dimensional vector space, and the 2-dimensional representation in UMAP does not always retain the distance relation in 16 dimensions.

**Fig 3.**
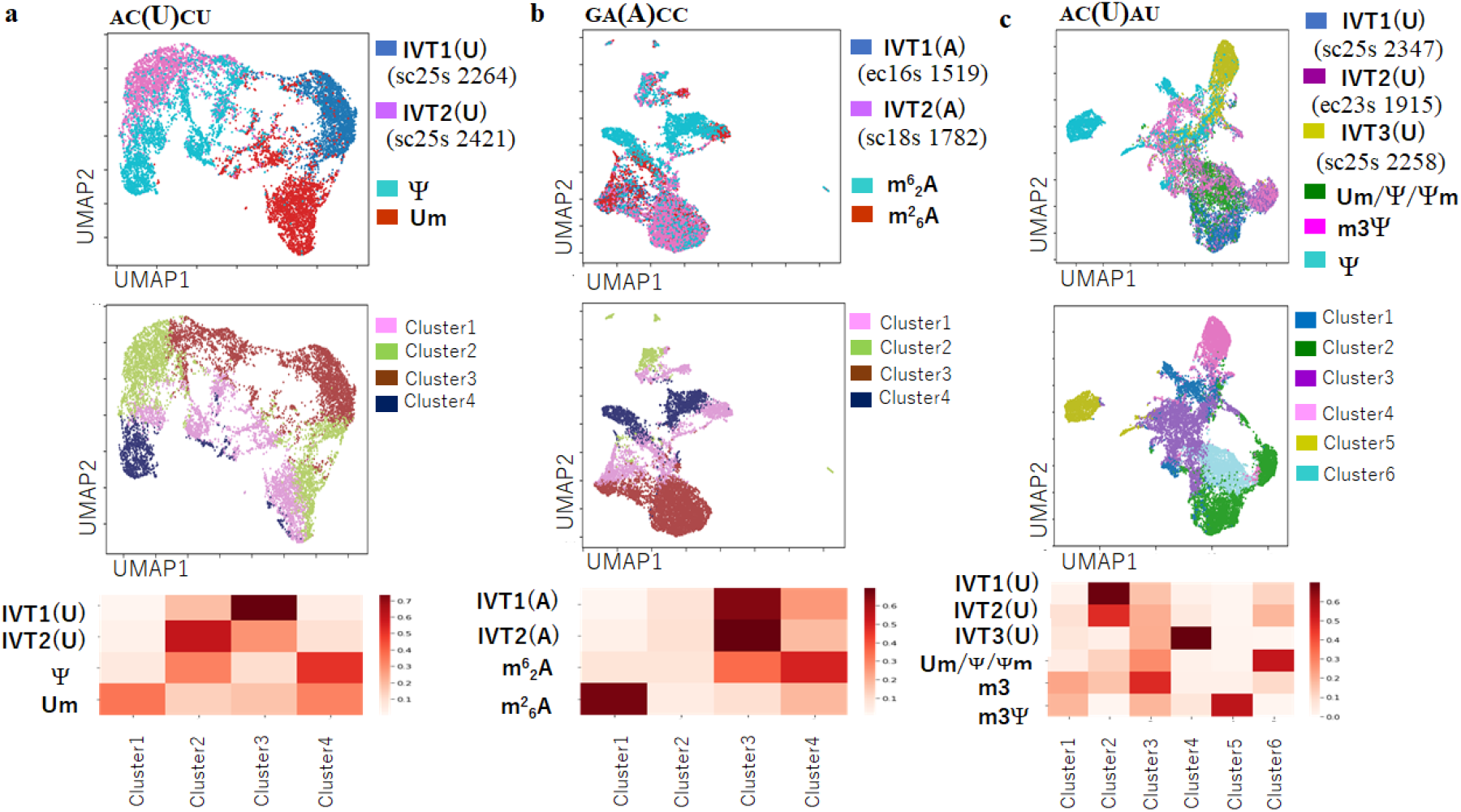
Separation of Modified signal. Separation of 5-mer sequence with multiple types of modified bases.(a~c) Upper row: UMAP representation of modified 5mer and unmodified IVT reference sequences. Middle row: K-mean clustering result mapped onto UMAP representation Bottom row: K-mean result for each labeled data set

Next, the overlap between the detected cluster and 5-mers of the modified and unmodified dataset was examined. The bottom row of the matrix shows how much of each data from different positions is assigned to each cluster. Four of seven cases, multiple IVT reference 5-mers from different position, clustered independently with more than 50% specificity (ex. a. ACUCU case); in the remaining cases, multiple reference 5-mer signals clustered in the same cluster with 50% or more specificity (ex. b. GAAAC case). There were 15 modified 5-mers in total, among which 10 modified 5-mers were assigned to an independent cluster with more than 50% specificity, even when the case corresponding to the reference IVT 5-mer was assigned in the same cluster. This result suggests that modified sequences are generally separated into unique clusters by their modification types (Fig 3 and S1 Table). In an exceptional case, one modified sequence was mainly assigned to the same cluster as the reference IVT, rather than forming a unique cluster, as observed for Am/m^6^2A and m4Cm/Cm, indicating the difficulty of separating either of these modifications from an unmodified signal. With the “ACUAU” 5-mer, I attempted to separate three modifications and corresponding IVT sequences (Fig 3c). For the *S. cerevisiae* position 2345, three modifications of Um/Ψ/Ψm have been previously reported [35]. However, the signal from this position hardly clustered with the Ψ modification cluster (position 2258), indicating that the Ψ modification at position 2345 in these data, if it exists, was very small.

### Analysis of a severe acute respiratory syndrome coronavirus 2 (SARS-CoV-2) dataset

I analyzed SARS-CoV-2 DRS data from three different groups in Korea [27], Australia [36], and England [37] using the procedure described above. Kim et al. prepared the IVT sequence, which was used as a reference to detect modifications. Ninety-one percent of reads were mapped to the reference in the IVT dataset, and 29-64% of native reads were mapped to the reference (S5 Table). A score draft with noise near the end of the IVT transcript was observed. To reduce error, I applied a window function for denoising. (S4 Method). The overall scores are shown in Fig 4a. All single nucleotide polymorphism sites in the English and Australian samples, against the Korean reference, were detected with scores from 0.05 to 1 (S6 Table). The other major peaks indicated the existence of PTMs. Many modifications have been previously reported in the 3’-terminal region centering on the N protein region (S10 Fig) [38][27]. I also conducted the LOGO [39] analysis with the nucleotide sequences of candidate sites and observed the weak trends of G/A/U enrichment at the −1 position of the target site and U/C enrichment in the +1 position of candidate sites. A score threshold of 0.04 was used, which corresponded to an expected false positive rate of 4~5 in (S7 Method) each candidate dataset, accounting for 101 candidates in total (S7 Table). More than half of the candidate sites from the three groups overlapped; however, 31, 16, and 11 candidates in the Korean, English, and Australian groups, respectively, remained as unique candidates (Fig 4b). In addition, 11 candidates overlapped with Kim et al. (Fig. 4c) with a match allowing for a ± 2 difference in the genomic position; however, most sites did not match because different software programs were used. I also compared the detected peaks to the short read RNA editing sites reported previously (S9 Fig) and observed almost no overlap between these two datasets. The RNA editing event, reported by a previous study that indicated a very low allele frequency (median variant allele frequency = 0.009) [40] on the SARS-CoV-2 transcriptome, may have been dismissed by this method. Further investigation, to identify each type of modification in the SARS-CoV-2 genome detected in this study, is ongoing.

**Fig 4.**
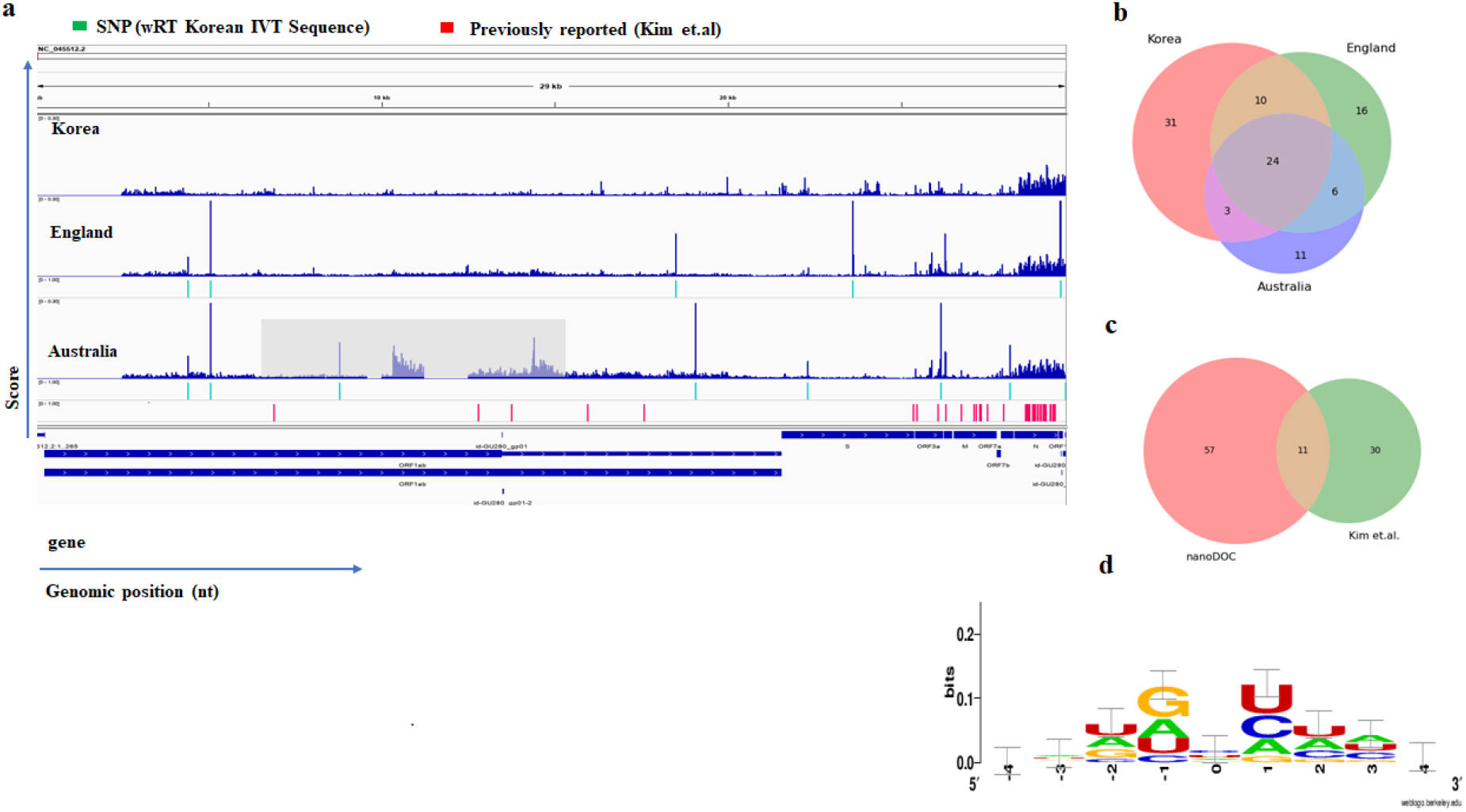
Modification analyses on SARS-Cov2 genome. a) Modification signal on SARS-Cov2 genome from Korea, Australia, and England the region covered by gray in Australia’s sample is the low depth region (under ×50), and hence it is not included in the final result. b) Intersection of modification candidates between the groups c) Intersection of modification candidates between nanoDoc and Tombo(Kim et. al) d) LOGO analysis of modification motif

### Availability and Future Directions

The source code and training weight used in this study have been deposited at https://github.com/uedaLabR/nanoDoc.

The future direction of developing this software is to accumulate data on different modifications for each 5-mer sequence and identify the different modification types. In addition, PTM detection without the IVT reference, as well as speed up and distributed commutation, are necessary to acquire an epitranscriptome map using this method for larger genomic size.

## Data Availability

The ribosomal RNA data were obtained from a previous study [20]. The INT/m6A modified sequence was also obtained from previous work [14], as were the SARS-CoV-2 DRS data for the three groups

[27],[37],[38]. Davidson et al. kindly shared the raw data from the DRS sample. The reference sequence was acquired from the National Center for Biotechnology Information (Bethesda, MD, USA) (S2 Table). The data supporting the findings of this study are available within the article [and/or] its supplementary materials.

## Acknowledgements

This work used the computational resources of AI Bridging Cloud Infrastructure (ABCI) provided by the National Institute of Advanced Industrial Science and Technology (AIST) through the High-Performance Computing Infrastructure (HPCI) research project (project ID: hp200145). A research and development grant from the Shimadzu Science Foundation (Kyoto, Japan) was applied to this study.

I would like to thank Dr. Hiroyuki Aburatani, Dr. Genta Nagae (The University of Tokyo, RCAST,Japan), Dr. Fumiko Kawasaki (RIKEN Center for Advanced Intelligence Project,Japan). and Dr. Shunpei Okada (Tokyo University of Science, Research Institute for Biomedical Science,Japan) for productive discussions and helpful advice. I would like to thank Dr David A. Matthews (University of Bristol, Department of Cellular and Molecular Medicine, UK), Dr. George Taiaroa (The University of Microbiology and Immunology, Australia), and Dr. Mark Akeson (UC Santa Cruz Genomics Institute & Biomolecular Engineering Department) for sharing data and assisting in locating specific fast5 files.

