## Supplemental_information for "nanoDoc: RNA modification detection using Nanopore raw reads with Deep One-Class Classification"

### Supporting Information

#### Supplemental methods

##### 1. Data preprocessing

Nanopore Multi-fast5/fast5 files containing raw electric signals were base-called with Guppy (v3.5.2; Oxford Nanopore Technologies [ONT], Oxford, UK), with the configuration file "res_rna2_r941_min_flipflop_v001.cfg" downloaded from ONT rerio project [23]. Using ont_fast5_apt [24], multi-fast5 reads were converted to single fast5. The Tombo (v1.5) [19] resquiggle command was used to map reads to their respective references (S2 Table) with the inner mappy/minimap2 aligner [25], and raw signals were further assigned to the mapped region of the reference (Fig 1a). The assigned raw signal for each genomic position was either up- or down-sampled to fit into an equal bin size of 60 (S12b Fig). The original signal length of each interval was recorded in other fields for use in a latter process. The apache parquet [26] format was used to record the data.

##### 2. Separation of training, test, and inference data

For initial training, 1200 reads are extracted from the curlcake unmodified IVT sample for 1024 5-mer, and 1,000 each (1024 × 1000) was used for training, with 200 used for validation. For DOC training, 10,000 reads were extracted from the curlcake unmodified IVT sample, and (1 target + 48 neighbor 5-mer × 10,000) reads were used for each 5-mer training. To avoid overfitting, for inference curlcake m6A modification, initial and DOC training was repeated with SARS-CoV2 *in vitro* samples (S4 Table). Although the curlcake sequence was designed to include all possible 5-mers in the sequence, the SARS-CoV2 genome is missing 64 5-mers of 1024 possible combination. Thus, for these missing combinations, the curlcake sample was used; however, 960 of 1024 (95%) 5-mers were from the SARS-CoV2 IVT sequence.

##### 3. Data formatting

The raw signal for each genomic position was either up- or down-sampled to fit into an equal bin size of 60.

For the IVT curlcake sample, 82.4% of the total reads were up-sampled, 0.4% reads remained the same, and 17.1% reads were down-sampled. The signal length for each segmented nucleotide showed a distribution with a median signal length of 23.0 and median absolute deviation of 23.72. A zero padding of size 10 was added to both sides of the data to form a 320 × 2 shape as an input.

#### 4. 5-mer in wider context

To determine how the 5-mer is selected from the actual read in wider context, the pre- and post-nucleotides of the 5-mer were evaluated in the *in vitro* curlcake sample [14] of 10 kb. Of the possible 16,384 7-mers, 7,366 combinations (46%) were found in the curlcake sample which was used as a training data set (S12c Fig). The training dataset was collected equally to contain the maximum number of combinations in the 7-mer context. Next, I examined whether the 7-mers in the training dataset caused a false-positive prediction.

The unmodified portion of rRNA dataset (first 500 nt in the reference of *E. coli* 23S and S. cerevisiae 25S rRNA) was used for this test. Between two cases, particularly if the 7-mer context was present in the training dataset, no significant difference and no false-positive value were observed, with all scores under 0.03; these values were below the detection threshold. The scores showed a mean of 0.0041 or the reference 7-mers and mean of 0.0039 without the reference 7-mers (S12d Fig).

#### 5. Choice of model and training

ADAM optimizers are used with a learning rate of 0.0005 and exhibit categorical cross-entropy loss. To evaluate the efficiency of the data format and model choice, three models were tested in addition to the nanoDoc model (model a, S13a Fig). The neural network model without dwell time information (with the same CNN architecture) (model b, S13b Fig), neural network model without the SENet architecture (model c, S13c Fig), and simple 8-layer CNN model, consisting of four layers of CNN (model d, S13d Fig) were tested. For model b, only the current channel was used as input; for model d, the raw signal intensity was averaged per aligned base. This test was performed to identify 1024 5-mer signal classifications using four models. The best validation accuracies with 400 epochs were a) 0.32, b) 0.31, c) 0.19, and d) 0.012, respectively.

#### 6. Scoring schema

The extracted feature of IVT1 was compared to those of the IVT2 and native signals, and the closest Euclidian distance in the target data set was calculated for each entry using approximate nearest neighbors. For the distance calculation, the 16-dimensional vector output from the CNN was used (S5a Fig). Each distance was summarized as a distribution; in the presence of a modified base, the distance was larger in the IVT1 vs. native comparison (S5b Fig). The distance distribution was plotted as a cumulative distribution function. The blue, orange, and green lines show the distance function from the IVT1 vs. IVT2, IVT1 vs. native, and native vs. IVT1 (reverse direction) comparisons, respectively. The surrounding area represents the disturbed signal feature from the IVT reference sequence. The score was calculated from this area and normalized by the sample size (S5c Fig).

#### 7. Adjustment and scoring

The Euclidian distance from the native signal to the IVT signal should be close or equal to that from IVT to IVT, if there are no modifications in the native sample; however, the value increases when modifications are present. Here, I defined cdf_IVT as the cumulative distribution function of the distance from IVT1 to IVT2 and cdf_Native as that from IVT1 to native. Across the range of distances, cdf_Native should be smaller than cdf_IVT in the presence of modified bases. For scoring, the area between two accumulated functions was calculated over the percentile threshold.

TB = Total number of data

x_thres = threshold x, where x is distance; 20 percentiles were calculated for each distribution of IVT 5-mers, and the threshold was set to that value.

Score = (Σ cdf_IVT(x) - cdf_Native(x))/TB, where x > x_thres.

#### 8. Depth evaluation

This analysis was performed using the rRNA dataset with varying depths of 10, 25, 50, 100, 250, 500, 1000, 2500, 5,000, 10,000, and 20,000 by randomly down-sampling the reads (S11a Fig). The AUC value decreases towards lower depths; however, the methods showed an AUC of 0.84 at a depth of 50 and AUC of 0.92 at a depth of 1000 (S11b Fig). The score was recalibrated so that all false-positive calls with scores of below 0.1 in all depths of 10–20,000 were suppressed.

#### 9. False-positive evaluation

To evaluate the false-positive ratio in an unmodified region, three consecutives unmodified 500-nt intervals in an rRNA sample were chosen, with varying depths of 50, 100, 250, 500, 1000, 2500, 5,000, 10,000, and 20,000 as described above for the down-sampling method. A total of 13,500 positions was examined (3 regions, 500 nt, 9 depth), producing a score distribution with a mean? and mad?. At a threshold of score 0.05, no false-positive candidates were detected above the threshold and two false-positives were detected at a threshold of 0.04 with a p-value of $1.4 \times 10^{-4e}$ (S8 Table). Applying this approach to the 29.9-kb SARS-CoV2 genome is expected to produce 4–5 (expect value of 4.56 ) false-positives genome-wide.

#### 10. ROC analysis

For all genomic positions, each position was labeled as either True or False. The known modification position in the rRNA dataset and ±1 positions were assigned as True and False elsewhere. To avoid an unequal degree of bias in ROC analysis, regions with a 50-nt margin, from the first modification site to the last modification site, were used.

#### 11. 5-mer modification analysis

The 5-mer signals were extracted from the native and corresponding reference IVT sequences. Each signal was subjected to a CNN, and the output signals were further analyzed using k-means clustering. Faiss [32] was used for k-means clustering, with a cluster size of 4–6 and 500 iterations. UMAP was used for visualization purposes, which reduced the dimensions from 16 to 2.

#### 12. Window function

The window function was applied for denoising.

For each point, the following functions were applied:

Let n be the arbitral genomic position and Vn be a score at position n,

$V_{baseline} = Max((V_{n-4}, V_{n-5})/2, (V_{n+4}, V_{n+5})/2)$ is to be used as the baseline

$V_{filter} = V_n - V_{baseline}$ (where, $n - 3 \leq x \leq n + 3$).

**S1 Fig. Data formatting**

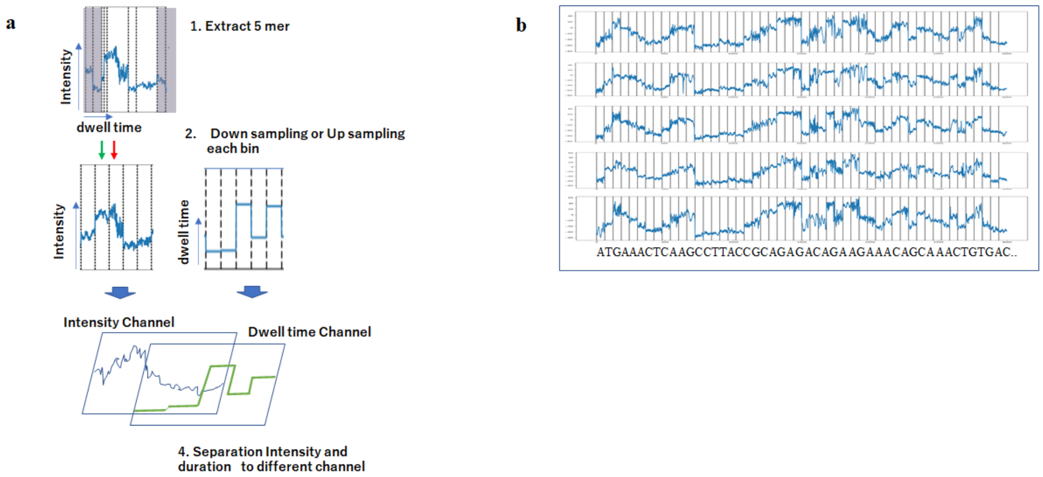


a) Schematic representation of signal uniformization and intensity-duration separation.

b) Representation of equaled bin sized signal.

**S2 Fig. Detailed architecture of 1D-CNN (modified from the study on Deep Binner [22] Fig 1).**

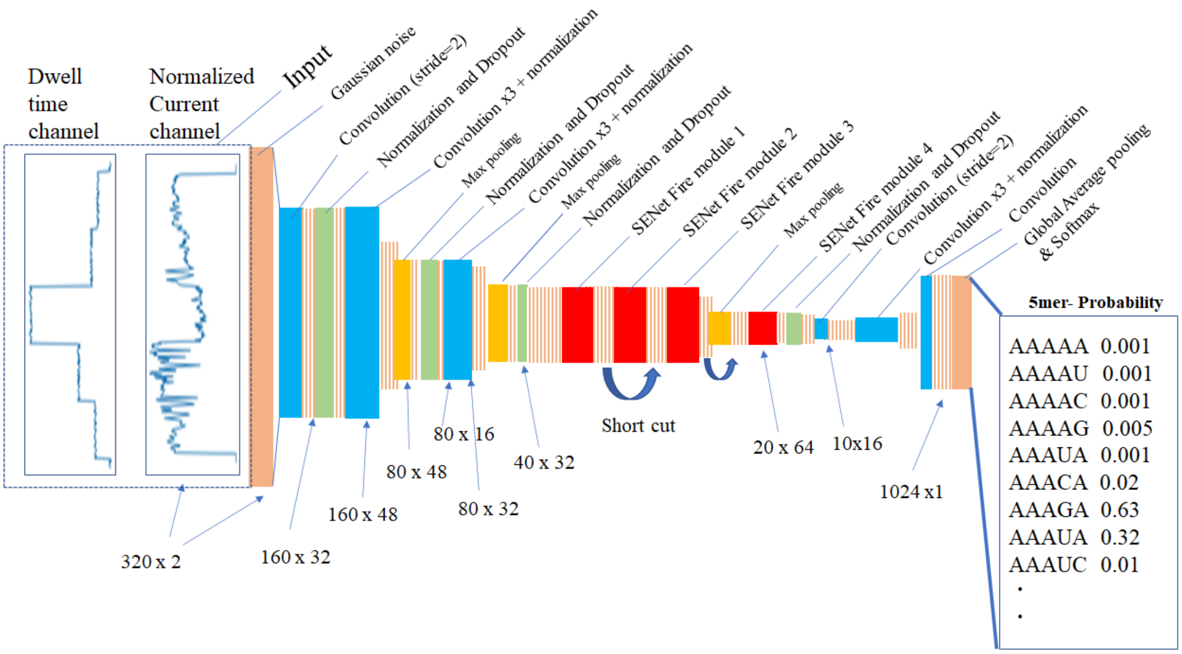

**S3 Fig . 1D-CNN training**

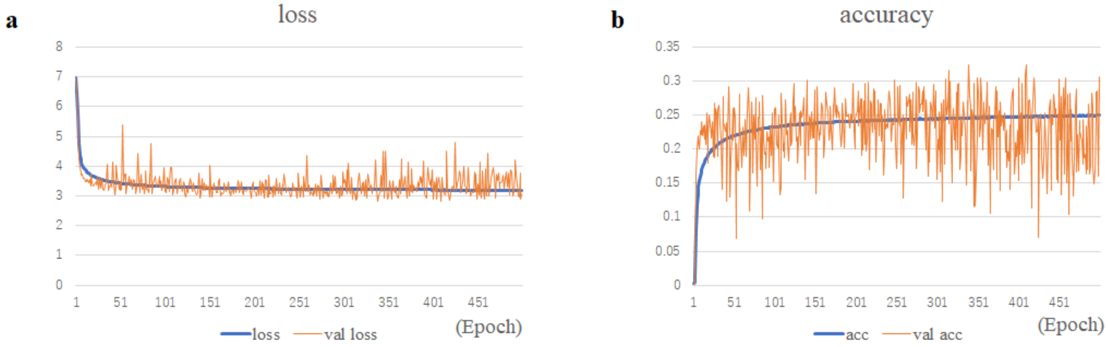


a) Loss of initial 1024 5-mer training of 1D-CNN. The blue line shows the loss values, and the orange line shows validation loss.

b) Accuracy of initial 1024 5-mer training of 1D-CNN. The blue line shows the accuracy values, and the orange line shows the validation accuracy.

**S4 Fig. Deep One-Class training**

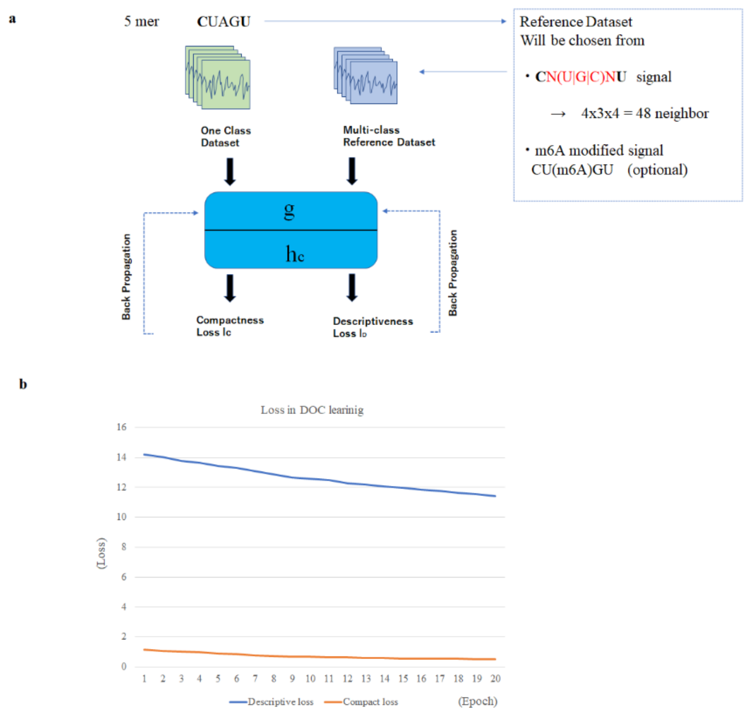


a) Target 5-mer dataset and reference dataset are input into the Deep One-Class model for training. Compactness loss is calculated and used in back propagation. Similarly, descriptive loss is used for reference network. The reference dataset is chosen with neighboring 5-mer where nucleotides at the 1$^{st}$ and 5$^{th}$ positions were same as in the target, and other nucleotide bases differed from the target 5-mer (figure is modified from the study on DOC [21] Fig 3).

b) Example of DOC training of “TCACG” 5-mer. Compactness loss and descriptive loss is plotted with training Epoch.

**S5 Fig. Scoring schema after feature extraction (figures obtained from the UCGAC vs UCG(m6A)C dataset).**

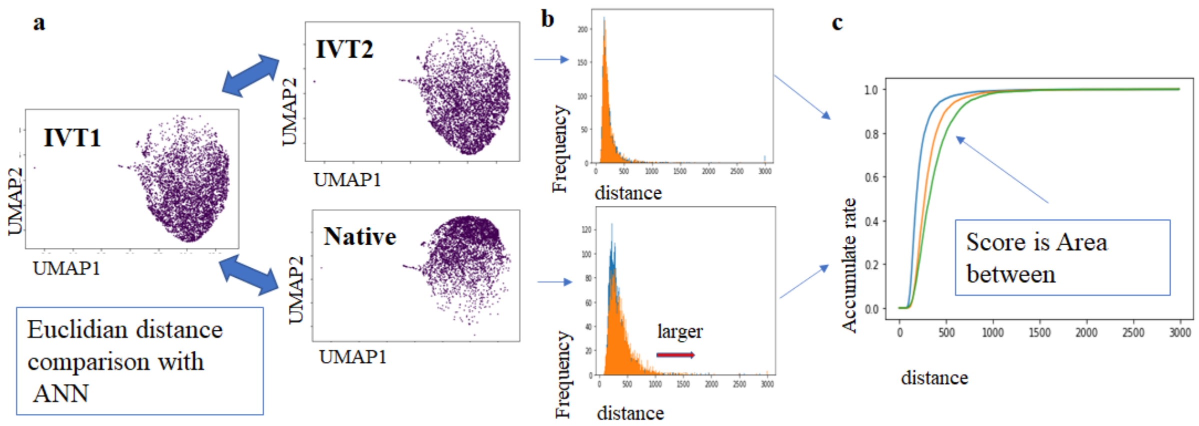


a) Extracted feature of IVT1 is compared to IVT2 and native signal, and the closest Euclidian distance in the target data set is calculated for each entry using approximate nearest neighbors. For the distance calculation, the 16-dimensional vector output from CNN is used.

b) Each distance is summarized as a distribution. When a modification is present, the distance is larger in the IVT1 vs native comparison.

c) Distance distribution is plotted as a cumulative distribution function. The blue line shows the distance function from IVT1 vs IVT2 comparison, the orange line shows IVT1 vs native comparison, and the green lines show native vs IVT1 (reverse direction).

Surrounded area reflects the disturbed signal feature from the IVT reference sequence. The score is calculated by normalizing this area to the sample size.

**S6 Fig. Result of m6A detection**

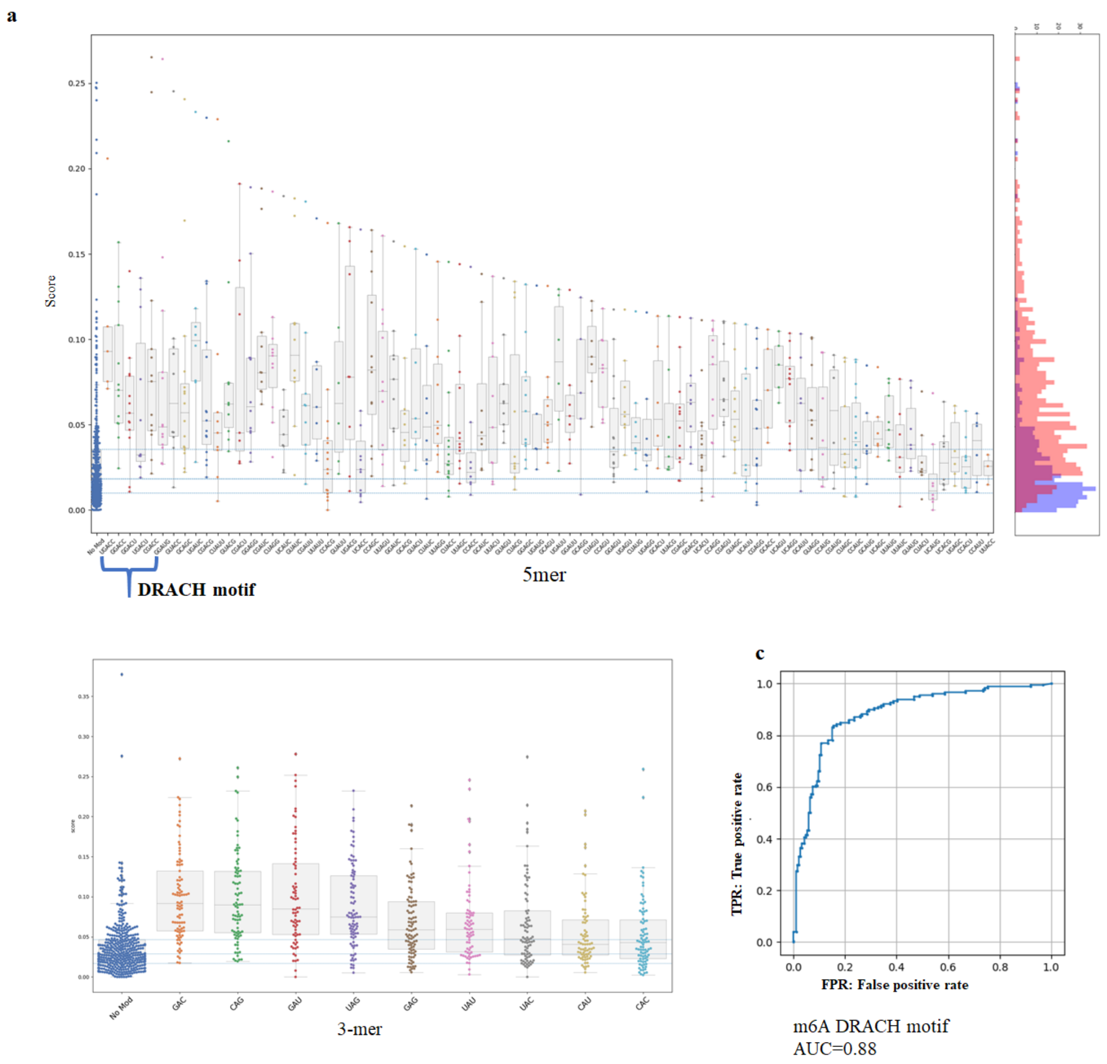


a) m6A score summarized by each 5-mer type and baseline with no modification (left most). Percentile line for baseline data was drawn for 75%, 50%, and 25% from the top. 5-mer with DRACH motif is shown on the left.

b) m6A score summarized by each 3-mer. G(m6A)C, C(m6A)G or G(m6A)U, and U(m6A)G are prone to detection.

c) ROC curve of m6A detection using artificially modified dataset. AUC values for nanoDoc, KSStats, and Tombo mean current difference were 0.96, 0.89, and 0.79, respectively.

**S7 Fig. Score vs modification ratio 62 modified positions reported previously with different read depths [34].**

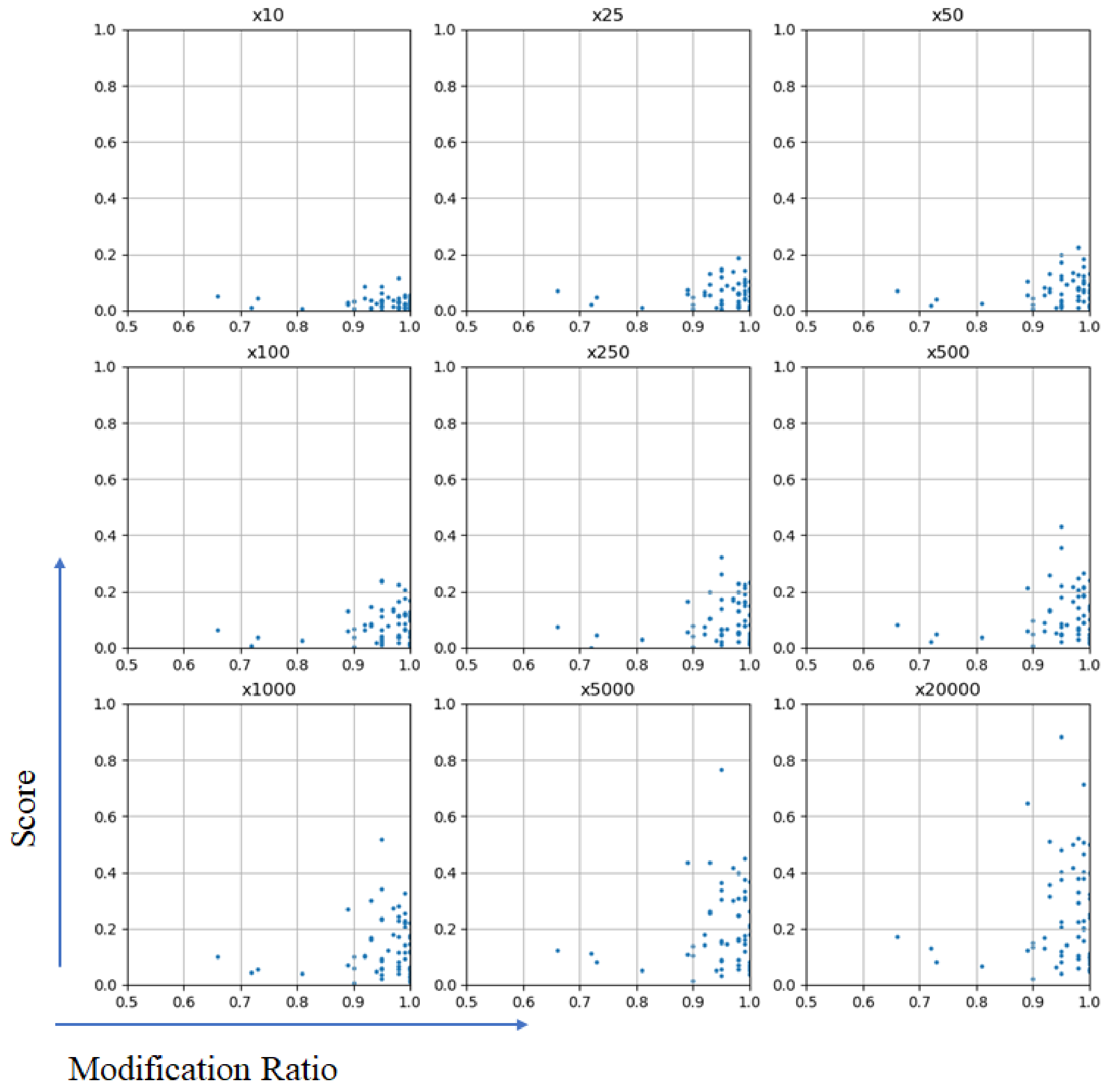

**S8 Fig. Score comparison of nanoDoc (upper panel) to KS statistics by Stephenson et al. (lower panel).**

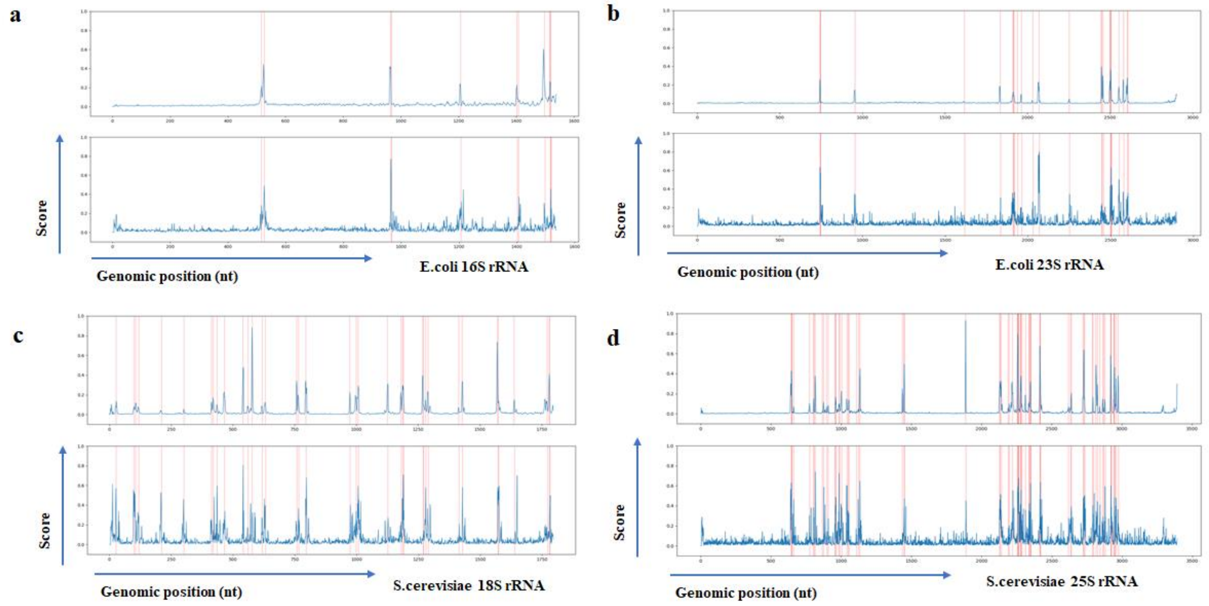


a) *E. coli* 16S rRNA, b) *E. coli* 23S rRNA, c) *S. cerevisiae* 18S rRNA, and d) *S. cerevisiae* 25S rRNA, respectively. The red line shows the position of known modification sites reported previously. The blue line shows the score level at each position.

**S9 Fig. Overlap with this study and single-nucleotide mismatch recurrently observed in short-read analysis reported earlier [40].**

**a**

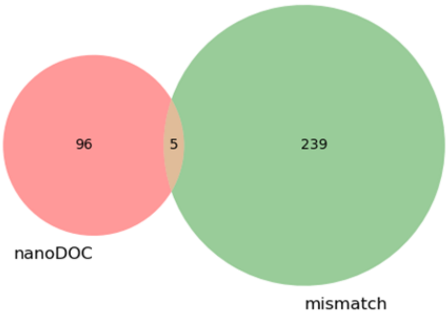


**b**

| Position | ref alt | allele variant frequency |
|---|---|---|
| 26132 | AG | 0.005 |
| 27913 | AG | 0.009 |
| 28549 | AG | 0.0089 |
| 29029 | TC | 0.0144 |
| 29280 | CT | 0.0166 |

**S10 Fig. RNA modification cluster on N protein region of the SARS-CoV2 genome.**

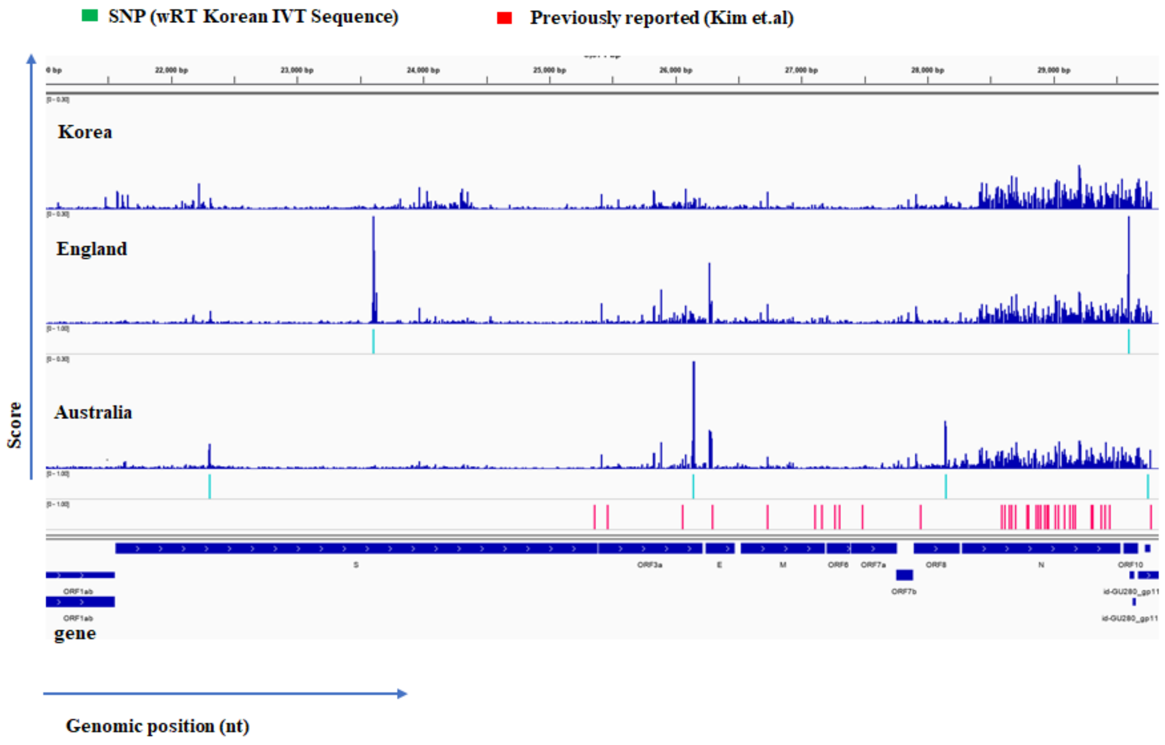

**S11 Fig. Depth evaluation of nanoDoc score**

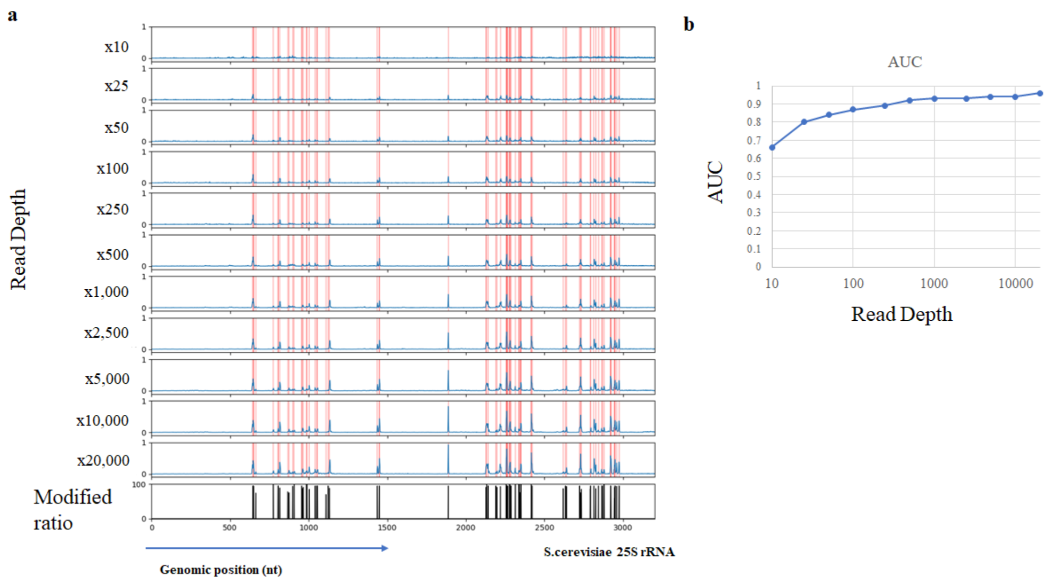


a) Score of *S. cerevisiae* 25S rRNA with varying read depths from 10 to 20,000. The red line shows the position of known modification sites in the literature. The blue line shows the score level at each position. The bottom graph depicts the modification ratio in the literature from earlier [34].

b) AUC value measured with different read depths using rRNA dataset.

**S12 Fig. Background analysis of data preprocessing**

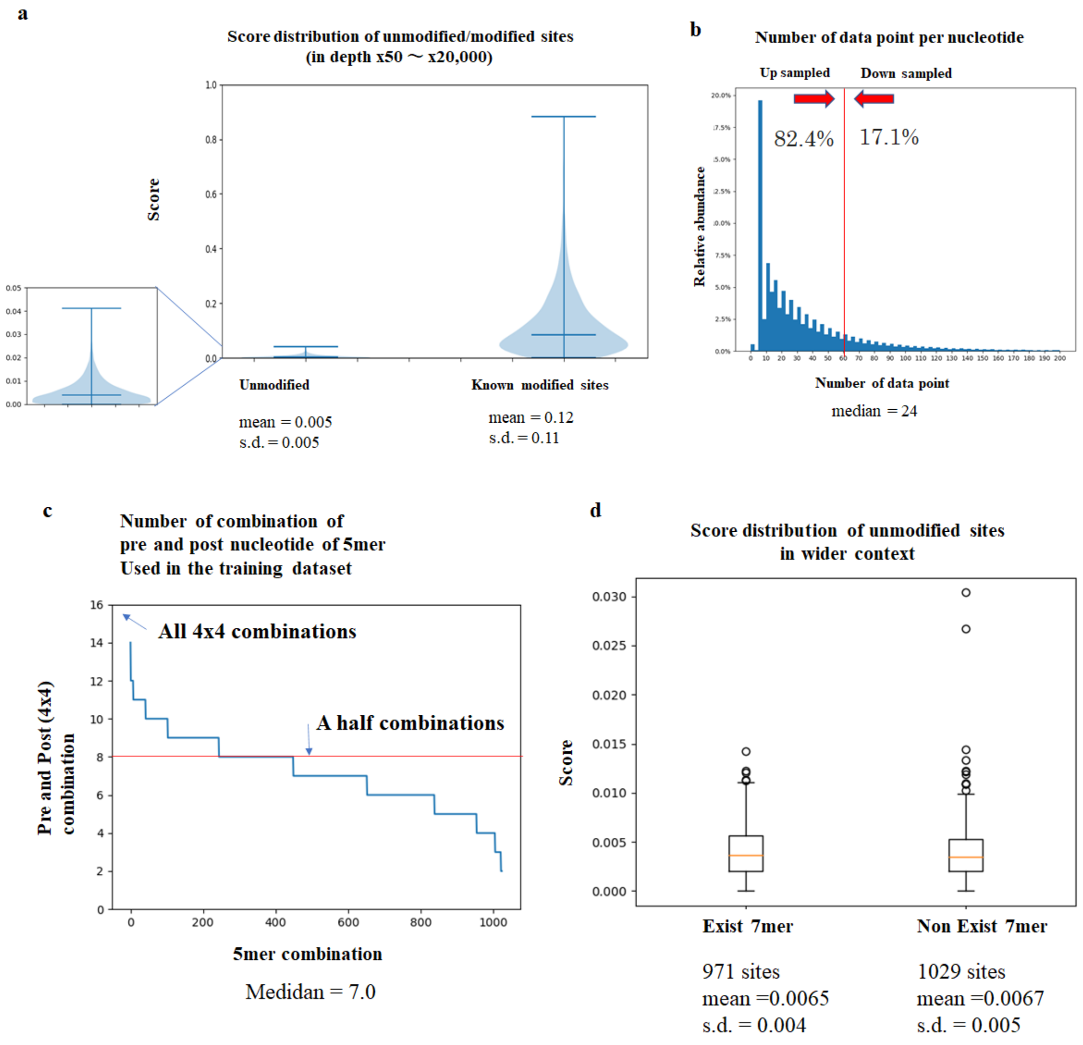


a) Score distribution of unmodified/modified sites. nanoDoc score evaluation with ×50 to ×20,000 depth using rRNA dataset. Unmodified sites were selected form four consecutive 500-nt intervals that do not contain known modifications.

b) Number of data points per nucleotide before formatting. Data point assignment to the nucleotide was performed using Tombo. For formatting, 82.4 samples were up-sampled to an interval length of 60, whereas 17.1% reads were down-sampled.

c) Number of combinations of pre- and post-nucleotide of 5-mer used in the training dataset.

d) Score distribution of unmodified sites in the two groups. Left box plot shows 7-mer context observed in the training dataset and right did not contain the 7-mer.

**S13 Fig. Another tested neural network models and data format.**

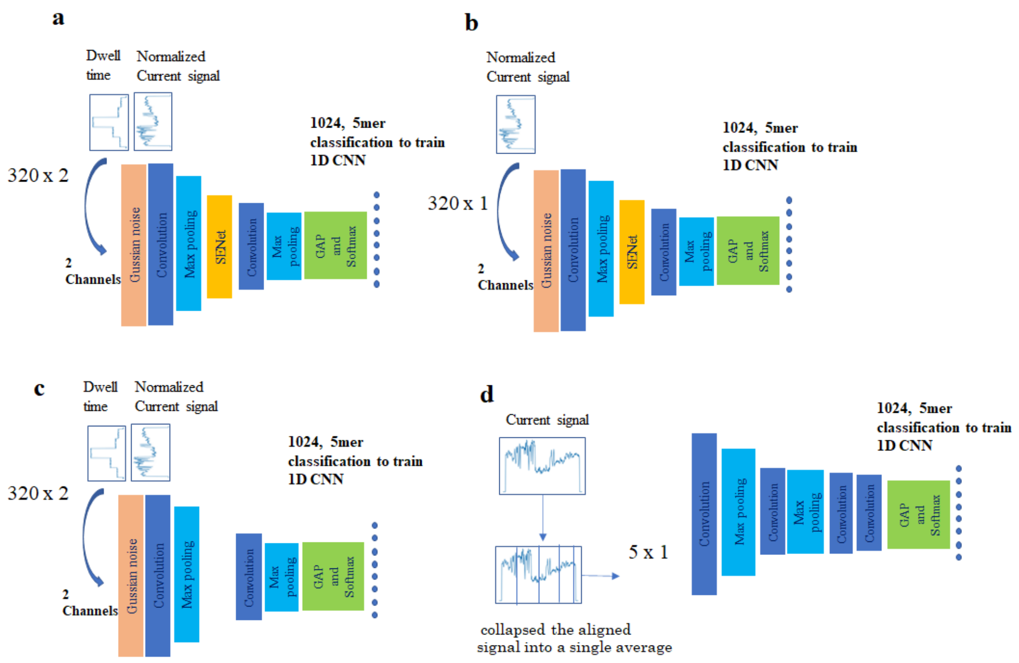


a) nanoDoc model as described in S2 Fig and above.

b) Same model as a. except that only the current channel is used as input.

c) Same model as a. except that the SENet layer is removed.

d) Six-layer CNN model where input dimension is 5 × 1, by collapsing the aligned signal into a single average value.

The best validation accuracy using 1024 5-mer is a) 0.32, b) 0.31, c) 0.19, and d) 0.012, respectively.

**S1 Table 5-mer with multiple modification and clustering result**

| 5mer | modifications | | |
|---|---|---|---|
| CGCCC | m4Cm | Cm | |
| | (ec16s 1402) | (sc18s 1639) | |
| CGUAA | m3U | Y | |
| | (ec16s 1498) | (ec23s 1911) | |
| GGAAC | m62A | Am | |
| | (ec16s 1518) | (sc18s 541) | |
| GAACC | m62A | m26A | |
| | (ec16s 1519) | (sc18s 1782) | |
| UUAAA | Am | m1A | |
| | (sc18s 619) | (sc25s 2142) | |
| ACUAU | Um/Y/Ym | m3Y | Y |
| | (sc25s 2347) | (ec23s 1915) | (sc25s 2258) |
| ACUCU | Y | Um | |
| | (sc25s 2264) | (sc25s 2421) | |

ec16s: *E.coli* 16S rRNA

ec23s: *E.coli* 23S rRNA

sc18s: *S.cerevisiae* 18S rRNA

sc25s: *S.cerevisiae* 25S rRNA

| CGCCC | cluster1 | cluster2 | cluster3 | cluster4 |
|---|---|---|---|---|
| IVT1 | 0.019 | 0.079 | 0.273333333 | 0.628667 |
| IVT2 | 0.135 | 0.049 | 0.404333333 | 0.411667 |
| m4Cm | 0.324 | 0.122333333 | 0.218333333 | 0.335333 |

| | | | | |
|---|---|---|---|---|
| Cm | 0.532 | 0.149 | 0.242333333 | 0.076667 |

| CGUAA | cluster1 | cluster2 | cluster3 | cluster4 |
|---|---|---|---|---|
| IVT1 | 0.023666667 | 0.189333333 | 0.328 | 0.459 |
| IVT2 | 0.160666667 | 0.554666667 | 0.015 | 0.269667 |
| m3U | 0.092333333 | 0.077333333 | 0.514333333 | 0.316 |
| Y | 0.265333333 | 0.175 | 0.012333333 | 0.547333 |

| CGCCC | cluster1 | cluster2 | cluster3 | cluster4 |
|---|---|---|---|---|
| IVT1 (ec16s 1402) | 0.019 | 0.079 | 0.273 | 0.629 |
| IVT2 (sc18s 1639) | 0.135 | 0.049 | 0.404 | 0.412 |
| m4Cm | 0.32 | 0.122 | 0.218 | 0.335 |
| Cm | 0.532 | 0.149 | 0.242 | 0.077 |

| CGUAA | cluster1 | cluster2 | cluster3 | cluster4 |
|---|---|---|---|---|
| IVT1 (ec16s 1498) | 0.024 | 0.189 | 0.328 | 0.459 |
| IVT2 (ec23s 1911) | 0.161 | 0.555 | 0.015 | 0.270 |
| m3U | 0.092 | 0.077 | 0.514 | 0.316 |
| Ψ | 0.265 | 0.175 | 0.012 | 0.547 |

| GGAAC | cluster1 | cluster2 | cluster3 | cluster4 |
|---|---|---|---|---|
| IVT1 | 0.087 | 0.600333333 | 0.043666667 | 0.269 |

| | | | | |
|---|---|---|---|---|
| IVT2 | 0.089666667 | 0.332333333 | 0.008333333 | 0.569667 |
| m62A | 0.097 | 0.324666667 | 0.048666667 | 0.529667 |
| Am | 0.134 | 0.002666667 | 0.836666667 | 0.026667 |

| GAACC | cluster1 | cluster2 | cluster3 | cluster4 |
|---|---|---|---|---|
| IVT1 | 0.008 | 0.082666667 | 0.654666667 | 0.254667 |
| IVT2 | 0.032 | 0.091666667 | 0.695 | 0.181333 |
| m62A | 0.076 | 0.076 | 0.350666667 | 0.497333 |
| m26A | 0.671666667 | 0.045666667 | 0.095666667 | 0.187 |

| UUAAA | cluster1 | cluster2 | cluster3 | cluster4 |
|---|---|---|---|---|
| IVT1 | 0.011333333 | 0.035333333 | 0.530666667 | 0.422667 |
| IVT2 | 0.018333333 | 0.698333333 | 0.083 | 0.200333 |
| Am | 0.011 | 0.02 | 0.250333333 | 0.718667 |
| m1A | 0.392666667 | 0.223 | 0.080333333 | 0.304 |

| ACUCU | cluster1 | cluster2 | cluster3 | cluster4 |
|---|---|---|---|---|
| IVT1 | 0.002333333 | 0.187666667 | 0.768 | 0.042 |
| IVT2 | 0.022666667 | 0.597 | 0.287333333 | 0.093 |
| Y | 0.057333333 | 0.331 | 0.105333333 | 0.506333 |
| Um | 0.359333333 | 0.140666667 | 0.173666667 | 0.326333 |

| ACUAU | cluster1 | cluster2 | cluster3 | cluster4 | cluster5 | cluster6 |
|---|---|---|---|---|---|---|
| IVT1 | 0.017666667 | 0.688666667 | 0.158666667 | 0.02 | 0 | 0.115 |
| IVT2 | 0.083666667 | 0.476666667 | 0.204333333 | 0.050667 | 0.002 | 0.182667 |
| IVT3 | 0.061 | 0.029333333 | 0.206 | 0.697667 | 0.005 | 0.001 |
| Um/Y/Ym | 0.036333333 | 0.121 | 0.277333333 | 0.018333 | 0.008 | 0.539 |
| m3Y | 0.221666667 | 0.157333333 | 0.479666667 | 0.02 | 0.02 | 0.101333 |
| Y | 0.187666667 | 0.002666667 | 0.187666667 | 0.064667 | 0.555333 | 0.002 |

**S2 Table. Reference file used in this study.**

*E. coli* (NC_000913.3.fa)

*S. cerevisiae* (R1-1-1_19960731.fsa)

Covid-19 sequences were downloaded from the GISAID (https://www.gisaid.org/) database.

Korea: SARS-COV2 isolate SNU01

England: SERS-COV2 BetaCoV/England/02/2020

Australia: SARS-COV2 Australia/VIC01/2020

Curlile transcript GEO database under the accession code GSE124309

**S3 Table.** AUC value measured using rRNA dataset with down-sampled reads depth.

| Depth | AUC |
|---|---|
| 10 | 0.66 |
| 25 | 0.8 |
| 50 | 0.84 |
| 100 | 0.87 |

| | |
|---|---|
| 250 | 0.89 |
| 500 | 0.92 |
| 1000 | 0.93 |
| 2500 | 0.93 |
| 5000 | 0.94 |
| 10000 | 0.94 |
| 20000 | 0.96 |

**S4 Table.** Train and test dataset used in this study

Table 4.1 Test and training dataset for rRNA, SARS-CoV2, and human transcript.

| | Initial training | DOC training | ref |
|---|---|---|---|
| | curlcake IVT | curlcake IVT | [14] |
| train | 1,000x1024 | 10,000x49 | |
| test | 200x1024 | 2,000x49 | |

Table 4.2 Test and training dataset for m6A detection

| | Initial training | DOC training | ref |
|---|---|---|---|
| | SARS-COV2 IVT,<br>curlcake IVT | SARS-COV2 IVT,<br>curlcake IVT | [27],[14] |
| train | 1,000x1024 | 10,000x49 | [ |
| test | 200x1024 | 2,000x49 | |

Table 4.3 Inference

| | | ref |
|---|---|---|
| rRNA | IVT | [20] |
| | WT | [20] |
| m6A | IVT | [14] |
| | WT | [14] |
| SARS-COV2 | IVT | [27] |
| | WT1 Korea | [27] |
| | WT2 England | [37] |
| | WT3 Australia | [36] |
| human | IVT | [10] |
| | WT | [10] |

**S5 Table. SARS-CoV2 DRS sequencing from three groups.**

| | Sequence ID | Virus | kit | size | Mapped reads /number of read | Tombo mapped % |
|---|---|---|---|---|---|---|
| Kim et al | MT039890.1 | SERS-COV2 isolate SNU01/ | SQK-RNA002<br>R9.4.1 | 730GB | 1,456,249 / 1,593,624 (in vitro)<br>570,916/879,679 (Infected) | 91.30%<br>64.90% |
| Taiaroa et al | MT007544.1 | SERS-COV2 Australia/VIC01/2020 | SQK-RNA002<br>R9.4.1 | 130GB | 198,765 / 680,347 (Infected) | 29.20% |
| Davidson et al | EPI_ISL_407073 (GISAID) | SERS-COV2 BetaCoV/England/02/2020 | SQK-RNA002<br>MIN106D R9<br>R9.4 | 160GB | 496,027 / 1,520,319<br>(Infected) | 32.60% |

**S6 Table. Score distribution at the position of SNP.**

| Position | England | Australia | ref | alt |
|---|---|---|---|---|
| 4402 | 0.15 | 0.03 | C | T |
| 5062 | 1 | 0.23 | T | G |

| | | | | |
|---|---|---|---|---|
| 8782 | - | 0.057 | T | C |
| 18488 | 0.13 | - | T | C |
| 19065 | - | 0.27 | T | C |
| 22303 | | 0.85 | T | G |
| 23605 | 0.73 | - | T | G |
| 26114 | - | 1 | G | T |
| 28144 | - | 0.19 | T | C |
| 29596 | 1 | - | A | G |
| 29750 | - | 0.55 | A | G |

**S7 Table. The candidate modification sites on the SARS-CoV2 genome detected using nanoDOC.**

| Genomic position | reference base | Korea(Score) | England(Score) | Australia(Score) |
|---|---|---|---|---|
| 4404 | A | 0.005 | 0.073 | 0 |
| 5064 | U | 0 | 0.334 | 0.159 |
| 12492 | C | 0.005 | 0.046 | 0 |
| 16325 | U | 0.057 | 0.027 | 0.009 |
| 16474 | G | 0.001 | 0.008 | 0.042 |
| 18486 | A | 0 | 0.044 | 0 |
| 19067 | A | 0 | 0 | 0.133 |
| 19986 | U | 0.075 | 0 | 0 |
| 21573 | U | 0.05 | 0.004 | 0 |
| 22222 | U | 0.072 | 0.003 | 0 |
| 23598 | U | 0 | 0.059 | 0.005 |
| 23607 | G | 0 | 0.393 | 0 |
| 23627 | G | 0 | 0.088 | 0.001 |

| | | | | |
|---|---|---|---|---|
| 23968 | U | 0.062 | 0.034 | 0.02 |
| 23969 | C | 0.058 | 0.045 | 0.016 |
| 24029 | U | 0.051 | 0.015 | 0.01 |
| 24296 | C | 0.046 | 0.002 | 0.003 |
| 24309 | C | 0.057 | 0.01 | 0.007 |
| 24351 | U | 0.049 | 0.013 | 0.01 |
| 25415 | C | 0.043 | 0.057 | 0.037 |
| 25830 | C | 0.052 | 0.047 | 0.043 |
| 25831 | U | 0.049 | 0.051 | 0.044 |
| 25887 | U | 0.018 | 0.096 | 0.075 |
| 26083 | U | 0.039 | 0.05 | 0.04 |
| 26084 | C | 0.056 | 0.032 | 0.036 |
| 26134 | C | 0 | 0 | 0.079 |
| 26146 | C | 0 | 0.016 | 0.539 |
| 26268 | A | 0 | 0.11 | 0.058 |
| 26271 | G | 0.003 | 0.169 | 0.109 |
| 26285 | U | 0 | 0.056 | 0.103 |
| 26287 | A | 0 | 0.064 | 0.052 |
| 26730 | U | 0.048 | 0.054 | 0.033 |
| 27910 | C | 0.043 | 0.046 | 0.032 |
| 27911 | U | 0.033 | 0.047 | 0.042 |
| 28307 | C | 0 | 0.008 | 0.042 |
| 28415 | A | 0.047 | 0.041 | 0.051 |
| 28429 | U | 0.073 | 0.055 | 0.058 |
| 28469 | U | 0.072 | 0.052 | 0.05 |
| 28494 | U | 0.04 | 0.028 | 0.029 |

| | | | | |
|---|---|---|---|---|
| 28541 | C | 0.041 | 0.044 | 0.034 |
| 28549 | C | 0.06 | 0.038 | 0.03 |
| 28562 | G | 0.063 | 0.041 | 0.049 |
| 28581 | U | 0.056 | 0.058 | 0.049 |
| 28596 | G | 0.053 | 0.036 | 0.05 |
| 28614 | A | 0.043 | 0.039 | 0.03 |
| 28644 | U | 0.064 | 0.05 | 0.045 |
| 28645 | G | 0.067 | 0.051 | 0.043 |
| 28659 | C | 0.044 | 0.024 | 0.022 |
| 28670 | U | 0.09 | 0.07 | 0.059 |
| 28671 | U | 0.092 | 0.075 | 0.058 |
| 28685 | C | 0.049 | 0.036 | 0.038 |
| 28704 | U | 0.088 | 0.082 | 0.075 |
| 28713 | C | 0.048 | 0.038 | 0.044 |
| 28764 | A | 0.045 | 0.036 | 0.021 |
| 28765 | A | 0.042 | 0.041 | 0.031 |
| 28803 | A | 0.048 | 0.051 | 0.05 |
| 28844 | G | 0.044 | 0.043 | 0.036 |
| 28845 | C | 0.039 | 0.041 | 0.044 |
| 28859 | A | 0.062 | 0.061 | 0.054 |
| 28916 | G | 0.061 | 0.042 | 0.035 |
| 28960 | C | 0.054 | 0.048 | 0.045 |
| 29017 | A | 0.074 | 0.081 | 0.064 |
| 29027 | C | 0.082 | 0.063 | 0.043 |
| 29044 | C | 0.079 | 0.064 | 0.076 |
| 29072 | C | 0.052 | 0.043 | 0.049 |

| | | | | |
|---|---|---|---|---|
| 29090 | C | 0.07 | 0.059 | 0.054 |
| 29100 | A | 0.049 | 0.035 | 0.029 |
| 29135 | A | 0.057 | 0.045 | 0.038 |
| 29169 | A | 0.044 | 0.032 | 0.021 |
| 29191 | U | 0.053 | 0.024 | 0.025 |
| 29202 | C | 0.123 | 0.09 | 0.077 |
| 29203 | G | 0.12 | 0.088 | 0.078 |
| 29211 | G | 0.115 | 0.085 | 0.078 |
| 29222 | U | 0.042 | 0.03 | 0.023 |
| 29265 | G | 0.071 | 0.053 | 0.053 |
| 29279 | C | 0.047 | 0.04 | 0.033 |
| 29300 | A | 0.062 | 0.059 | 0.059 |
| 29301 | U | 0.061 | 0.061 | 0.054 |
| 29315 | A | 0.045 | 0.046 | 0.041 |
| 29374 | C | 0.066 | 0.051 | 0.045 |
| 29414 | C | 0.074 | 0.054 | 0.07 |
| 29415 | C | 0.073 | 0.057 | 0.075 |
| 29427 | A | 0.043 | 0.035 | 0.024 |
| 29462 | C | 0.042 | 0.021 | 0.022 |
| 29477 | A | 0.049 | 0.042 | 0.06 |
| 29493 | G | 0.041 | 0.029 | 0.021 |
| 29510 | G | 0.069 | 0.051 | 0.047 |
| 29543 | A | 0.085 | 0.072 | 0.061 |
| 29572 | G | 0.053 | 0.041 | 0.045 |
| 29594 | U | 0.03 | 0.137 | 0.019 |
| 29598 | U | 0 | 0.106 | 0 |

| | | | | |
|---|---|---|---|---|
| 29602 | C | 0.054 | 0 | 0.037 |
| 29654 | U | 0.057 | 0.039 | 0.035 |
| 29664 | U | 0.074 | 0.052 | 0.057 |
| 29676 | A | 0.084 | 0.069 | 0.058 |
| 29687 | G | 0.061 | 0.043 | 0.031 |
| 29688 | U | 0.059 | 0.044 | 0.029 |
| 29736 | G | 0.074 | 0.05 | 0.008 |
| 29737 | C | 0.078 | 0.05 | 0.013 |
| 29769 | A | 0 | 0 | 0.052 |
| 29773 | C | 0.048 | 0.035 | 0 |

**S8 Table. Score distribution of unmodified bases in test sample and the corresponding threshold for detection.**

| Score threshold | number of unmodified sites above threshold | P-value |
|---|---|---|
| 0.05 | 0 | 0 |
| 0.04 | 2 | 0.000148 |
| 0.03 | 30 | 0.002222 |
| 0.02 | 274 | 0.020296 |
| 0.01 | 2026 | 0.150074 |
| 0 | 13500 | |
